# Standard metabolic rate does not associate with age-at-maturity genotype in juvenile Atlantic salmon

**DOI:** 10.1101/2021.08.25.457673

**Authors:** Eirik R. Åsheim, Jenni M. Prokkola, Sergey Morozov, Tutku Aykanat, Craig R Primmer

## Abstract

Atlantic salmon (*Salmo salar*) is a species with diverse life-history strategies, to which the timing of maturation contributes considerably. Recently, the genome region including the gene *vgll3* has gained attention as a locus with a large effect on salmon maturation timing, and recent studies on the *vgll3* locus in salmon have indicated that its effect might be mediated through body condition and accumulation of adipose tissue. However, the cellular and physiological pathways leading from *vgll3* genotype to phenotype are still unknown. Standard metabolic rate is a potentially important trait for resource acquisition and assimilation and we hypothesized that this trait, being a proxy for the maintenance energy expenditure of an individual, could be an important link in the pathway from *vgll3* genotype to maturation-timing phenotype. As a first step to studying links between *vgll3* and the metabolic phenotype of Atlantic salmon, we measured the standard metabolic rate of 150 first year Atlantic salmon juveniles of both sexes, originating from 14 different families with either late maturing or early maturing *vgll3* genotypes. No significant difference in mass-adjusted standard metabolic rate was detected between individuals with different *vgll3* genotypes, indicating that juvenile salmon of different *vgll3* genotypes have similar maintenance energy requirements in the experimental conditions used and that the effects of *vgll3* on body condition and maturation are not strongly related to maintenance energy expenditure in either sex at this life stage.

**Summary statement:** We show that *vgll3*, a gene known to have significant effects on Atlantic salmon (*Salmo salar)* life-history strategy, does not associate with standard metabolic rate in salmon juveniles.

## INTRODUCTION

Due to their close links with fitness, life-history traits such as age at maturity, offspring number and size have been a long-term focus of research in biology (Stearns, 1989). Recent advances in genomics have enabled the identification of genes associated with such traits in an increasingly broad range of species (e.g. Barson et al., 2015; Lamichhaney et al., 2016; Narum et al., 2018; Troth et al., 2018). However, follow-up studies for determining the mechanistic basis underlying such associations remain rare, and so, a thorough understanding of the underlying biology as well as the wider consequences of these genes is lacking.

Age at maturity is a key life-history trait as it is the source of a trade-off between current and future reproduction (Roff, 2002; Stearns, 1989), and the different life-history strategies of the Atlantic salmon (*Salmo salar*, Linnaeus, 1758) are a good example of this trade-off. Atlantic salmon spend a significant part of their life at sea to grow before they return as sexually mature fish, often in their home river, and there is significant variation in the time spent at sea (sea age at maturity). This time varies from one to five years, with an individual’s size roughly doubling with every extra year spent feeding at sea (Hutchings & Jones, 1998). Spending more time at sea gives a potential fitness advantage, as larger, later-maturing individuals have a higher reproductive success (Mobley et al., 2020). However, the increased time at sea also comes with a higher risk of dying before first reproduction, resulting in a trade-off between size at maturity and risk of mortality (Stearns, 2000).

Analyzing the genetic background for this variation, a genome-wide association study (GWAS) covering 57 European Atlantic salmon populations identified a large-effect locus on chromosome 25 that is significantly associated with age at maturity, explaining 39% of the variation in this trait (Barson et al., 2015). The strongest candidate gene in this region was the vestigial-like family member 3 (*vgll3*) gene. The two alleles of this locus, *E* and *L*, associated with either early or late age at maturity, respectively, as well as shorter or longer body length (in equally aged returning individuals). Subsequently, multiple studies have confirmed the large effect of this locus. (Ayllon et al., 2015; Debes et al., 2021; Verta et al., 2020).

Taking studies of this gene to the lab, examining body condition and maturation probability in a common garden setting, Debes et al. (2021) found that the *vgll3*\**E* allele associated with a higher body condition in males and females (but not body length), as well as a higher probability of male maturation in the first autumn after hatching. Interestingly, Debes et al. (2021) also showed that maturation probability in males could be predicted using the body condition of their sisters (all immature), demonstrating a connection between *vgll3* and body condition that is not caused by the enlarging gonads in maturing individuals. Additionally, expression of *vgll3* has been linked with activation of the intracellular HIPPO signalling pathway, suggesting a role in inhibiting gonad development and promoting adipocyte differentiation (Kjærner-Semb et al., 2018; Kurko et al., 2020; Verta et al., 2020). Thus, the findings on the mechanistic basis of the *vgll3-*maturation pathway so far indicate that different *vgll3* genotypes are causing differing patterns of resource allocation or assimilation, where the *vgll3*\**E* allele is positively influencing juvenile body condition, enabling earlier gonadal development in males. However, many aspects of the complete molecular and physiological pathways for this process remain unclear.

Early-life performance in acquiring and efficiently using resources sets the stage for the timing of maturation: sexual maturation and reproduction are costly processes, and a key part of juvenile development is the accumulation and storage of excess energy to support future maturation (Hutchings & Jones, 1998; Simpson, 1992; Thorpe, 2007). Standard metabolic rate (SMR)(Chabot et al., 2016) is a potentially important trait with regards to this as it relates to the organism’s maintenance energy requirements, capacity for nutrient assimilation, and responses to changing resource availability (Armstrong et al., 2011; Auer, Bassar, et al., 2020; Auer, Solowey, et al., 2020; Bochdansky et al., 2005; Millidine et al., 2009; Rosenfeld et al., 2015). The effect of *vgll3* genotype on body condition, as well as its strong effect on age at maturity, raises the question of whether *vgll3* might assert some of its effects via SMR, warranting an investigation on the potential influence of *vgll3* on metabolic phenotype. Additionally, beyond the effects on growth, body condition and gonad development, little is known about the effects of *vgll3* on a broader range of physiological systems. Given the range of traits found to associate with SMR, testing for an association between SMR and *vgll3* could then help in refining the potential traits and physiological processes in the scope of *vgll3*’s effects, such as differences in digestive capacity (Millidine et al., 2009), enzyme activity (Norin & Malte, 2012), mitochondrial leak respiration (Salin et al., 2016, 2019), and behaviour (Binder et al., 2016; Biro & Stamps, 2010; Cutts et al., 1998; Metcalfe et al., 1995; Yamamoto et al., 1998)

To further our understanding of the mechanistic functions of *vgll3*, in this study we test if there is an association between SMR and *vgll3* genotype in a common-garden environment using 150 juvenile Atlantic salmon (69 males and 81 females) from 14 families with either *vgll3*\**EE* or *vgll3*\**LL* genotypes. Based on 1) earlier studies indicating *vgll3* as a resource allocation locus and 2) the importance of metabolic rate for resource acquisition toward growth, we hypothesised that *vgll3* genotype is influencing energy budget allocation via an effect on physiological traits relating to metabolism.

## MATERIALS AND METHODS

### Experimental animals and husbandry

The experiment was conducted under an animal experiment permit granted by the Finnish Project Authorisation Board (permit nr. ESAVI/4511/2020). The cohort of Atlantic salmon used in this study was established using parental individuals deriving from a first-generation hatchery broodstock of salmon originating from the river Kymijoki in Finland, managed by the Natural Resources Institute Finland (LUKE) at their hatchery in Laukaa. Eggs and milt were collected and transferred to the experimental facilities at the University of Helsinki in October 2019 and parents were crossed to create 14 full-sib families (seven *vgll3*\**EE* families and seven *vgll3*\**LL* families). Fertilized eggs were incubated in darkness in vertical incubators in replicated, family-specific compartments, with a water temperature of 7°C until March 2020.

After hatching, on March 6^th^, 2020, the alevins were transferred to the experimental facilities at Lammi Biological Station (61°04′45′′N, 025°00′40′′E, Lammi, Finland), several weeks before they commenced independent feeding. Each family was reared in a randomly selected separate circular 165-litre tank (90cm diameter) and supplied with a continuous flow of UV filtered water from the local lake Pääjärvi, warmed by 1°C through a heat-exchange system. The incoming water was directed to create a slow circular flow in the tank. The photoperiod was adjusted according to the local latitude for the entire experiment. Initially, fish were fed eight times/day with commercial 0.2 mm pellet food (Vita, Veronesi, Italy), i.e. ad libitum. As fish grew, the 0.2 mm feed was gradually replaced with 0.5 mm pellet food at an increasing frequency (up to 12 times/day). The temperature rose gradually from an average of 4.7°C in March to 11.5°C in July (Fig. 1), and the average temperature during this period was 7.38°C. Tanks were flushed of uneaten food daily, and tank surfaces were carefully scrubbed clean of dirt and algae once or twice each week (depending on temperature). Tanks were checked for dead individuals daily; The mortality rate throughout the study was 3.9%. The specific *vgll3* genotype of each family was not known to people participating in fish husbandry, nor to those conducting the respirometry experiments, and was only revealed after the respirometry data had been finalised.

**Figure 1.**
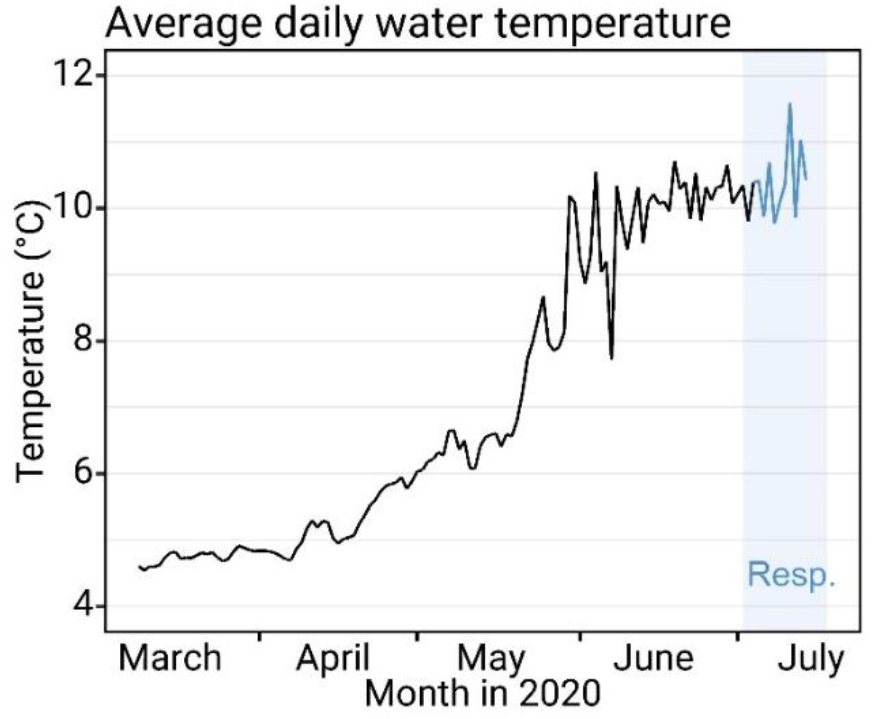
Average daily water temperature in holding tanks at Lammi Biological Station. Water was supplied from the local lake Pääjärvi, and temperature thus fluctuated according to the temperature of the lake. Average temperature of the holding tanks during the respirometry period and the 3 days leading up to it was 10.34±0.77°C (SD). All tested fish were held at a constant temperature of 10.5°C in acclimation tanks for 40 hours before respirometry.

### Overall respirometry procedure

Starting on July 4^th^, 2020 (at about 1900 degree days), one batch of 16 fish was tested using 16 respirometers (Appendix 1 - Fig. A1) each day for 10 days (except on July 5^th^, and with one gap day after 5 days for cleaning and sensor calibration), resulting in a total of 160 individuals in the experiment. Fish from the 14 families were evenly distributed among the 16 respirometers for each batch, along with extra individuals from two families, rotating the extra families between batches (Fig. 2). Prior to respirometry trials, fish were held in an acclimation tank with a constant water temperature of 10.5°C without feed for 40h. To maintain the family identity of the fish in the acclimation tank, the fish were individually kept in small 20×20×10cm cages (stainless steel cage lined with a plastic mosquito-net mesh) submerged in the acclimation tank. Air exposure of fish was avoided both during capture from holding tanks and when moving them to respirometers by transferring fish in small plastics cups filled with water. A detailed description of the respirometry holding system is in appendix 1.

**Figure 2.**
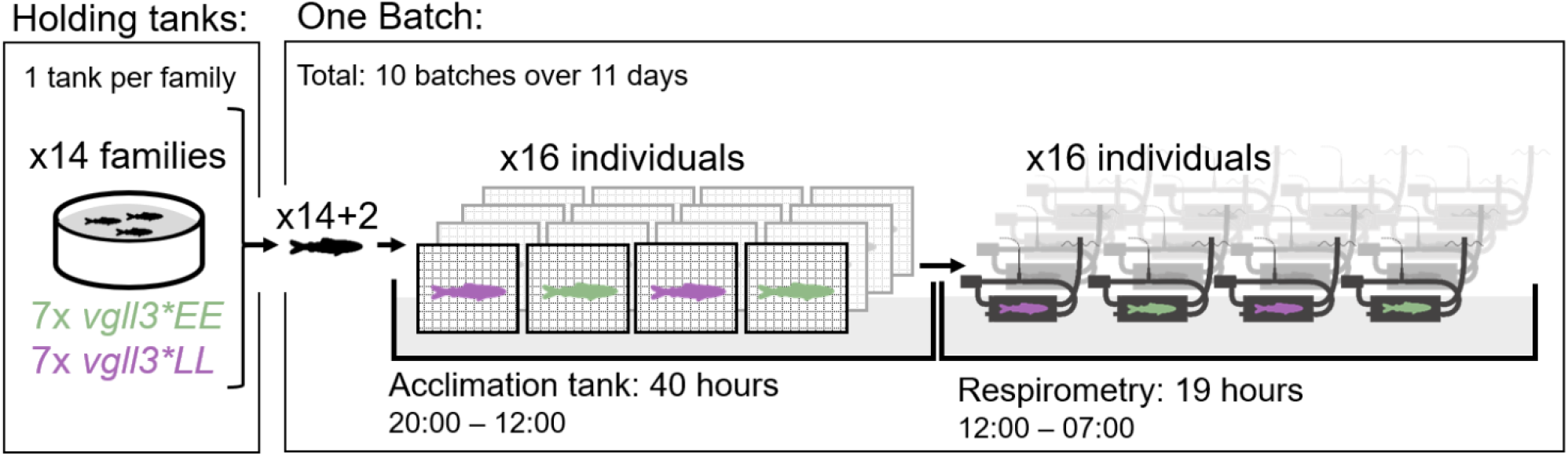
Overall respirometry procedure and structure. Fish were reared in 14 tanks, separated by family, with each family being either *vgll3*\**EE* or *vgll3*\**LL* genotype. Fish were tested in 10 batches of 16, with an acclimation period of 40 hours (constant temperature, no feed) before being held in respirometers for about 19 hours while oxygen consumption was being measured.

### Respirometry details

Standard metabolic rate was measured in fish at rest using intermittent flow respirometry (Chabot et al., 2016; Forstner, 1983; Steffensen et al., 1984). A detailed description of the SMR definition, protocol and respirometry system is in appendix 1 – supplementary materials and methods. Fish were put in their respirometers around 12:00 at noon, and the procedure was stopped at 7:00 in the morning the next day (total time 19h). Temperature was kept the same as in the acclimation tanks at 10.53±0.02 °C (mean±SD) during respirometry. The respirometry tank was covered with a light-blocking tarp to leave the respirometers in complete darkness during the entire procedure. The respirometry system was in a separate room from other activities in the research building to avoid disturbances.

Respirometers were set to cycle between the open flush phase and the closed measurement phase in 5-(flush) and 15-minute (measurement) intervals, resulting in a total of 50-60 measurement cycles for each batch. Sampling frequency of oxygen concentration was set to one sample every two seconds. The average minimum oxygen concentration reached during the closed phase was 7.58±0.07 mg L^-1^ (±SE). For the entire experiment, the lowest overall oxygen concentration reached was 4.22 mg L^-1^, and there were ten cycles in total where the oxygen concentration reached below 6 mg L^-1^.

The insides of respirometers were brushed daily to minimize bacterial or algal growth, and oxygen probes were gently wiped with a paper tissue moistened with a 70% ethanol solution before the measurements. After half of the batches were completed, the oxygen probes were recalibrated and the entire system (except acclimation tanks) was cleaned and disinfected with a solution of bleach, followed by thorough rinsing with fresh water. Background respiration was recorded for each batch of measurements before and after putting the fish in the respirometer. This was done individually for each chamber in each batch. The length of each background respiration measurement was 15 minutes (one closed cycle). The average background respiration was 7.48±4.55 % (±SD) of the total respiration in the chamber with fish.

After each respirometry batch, fish were removed from their respirometers and euthanized (sodium bicarbonate-buffered methanesulfonate overdose, 250 mg L^-1^), then carefully dried on each side using a tissue paper before their body mass was measured to the nearest 0.01gm using a precision scale (Scout STX222, Ohaus, Parsippany, USA). A fin clip was taken from the caudal fin and stored in ethanol for verifying the *vgll3* genotype and for sex determination. We used Kompetitive allele-specific polymerase chain reaction (KASP^*TM*^, LGC, UK) assays (He et al., 2014) for the *vgll3*_*TOP*_ SNP and the sex-specific SDY locus as described in (Sinclair-Waters et al., 2021).

### Statistical analysis

All data analysis was performed in the *Rstudio* v. 1.4.1717 (RStudio Team, 2020) software environment running *R* v. 4.0.4 (R Core Team, 2021). R packages used for analysis were *FishResp* v. 1.1.0 (Morozov et al., 2020) for extraction of respirometry data; *lme4* v. 1.1.26 (Bates et al., 2021, p. 4) and *lmerTest* v.3.1.3 (Kuznetsova et al., 2020) for mixed models analysis; *ggplot2* v. 3.3.5 (Wickham et al., 2021, p. 2) for data visualisation; *ggeffects* v. 1.1.0 (Lüdecke et al., 2021) for prediction of marginal means; *cvms* v 1.3.1(Olsen et al., 2021) for performing cross-validation of models; and *mclust* v. 5.4.7 (Fraley et al., 2020) for calculating the mean lowest normal distributions used for estimation of SMR. A detailed description of the calculation of SMR from the respirometry data is in appendix 1 – supplementary materials and methods.

At the time of respirometry testing, the body mass distributions of the *vgll3*\*EE and *vgll3*\*LL families were slightly different. This was caused by an earlier experiment utilizing the same study population, which sampled individuals non-randomly in an attempt to size-match individuals from different families, causing significant body mass difference between *vgll3* genotypes in this experiment that is not necessarily caused by the *vgll3* genotype. As a consequence of this, no statistical analysis on body mass differences was done, and all analyses controlled for body mass.

To test if *vgll3* genotype associated with a change in standard metabolic rate, we used a linear mixed model where log_10_ SMR (mg O_2_ L^-1^ h^-1^) was fitted against log_10_ body mass (g) (accounting for allometric scaling), including *vgll3* genotype (LL/EE), sex (m/f) and the interaction between *vgll3* genotype and sex as fixed effects, thus including both the direct effects of *vgll3* genotype and sex on standard metabolic rate, as well as a potential sex difference in the effect of *vgll3*. Family (1-14) was included in the model as a random effect, accounting for variation caused by both tank- and family effects as they were confounded in the experiment (one family per tank). Respirometry batch (1-10) was also included as a random effect. To control for potential temperature effects on metabolic rate via thermal acclimation, the average holding tank temperature (°C) for the last three days before the fish entered the acclimation tank was included as an additional fixed effect.

To test if there was an association between *vgll3* and body condition, we used a similar approach as above, where we modelled body condition in a mixed effect model as a function of *vgll3* genotype, sex and the interaction between *vgll3* genotype and sex as fixed effects, as well as family and batch as random effects. Body condition was calculated as Fulton’s K, which is the body weight (g) divided by the cube of the body length (mm) multiplied by 100, and represents the relationship between the body weight and length (Ricker, 1975).

p- and F-test values were calculated for fixed effects using Satterthwaite’s method with type III tests in models with interactions, and type II tests in models without interactions. The full models were compared to two alternative models: A sex-only model removed *vgll3* genotype and its interaction with sex as fixed factors, and a simple model removed both sex and *vgll3* genotype. These models were compared to the full model using Akaike’s information criteria (AIC) and by testing the predictive power of each model by cross-validation. Cross-validation was performed by separating the data set into two sets of either even or odd-numbered respirometry batches, and then using the model parameters derived from each set try predicting the SMR of the other; The Root-mean-square error was then used to compare the predicted values against the observed ones for each model. All models were visually examined and confirmed to have normally distributed residuals. Statistical summaries of both sets of models are shown in Table A1 and A2 in appendix 2 – supplementary analysis and data.

Nine individuals were removed from the analysis because of technical issues (pump, oxygen logger, or identification failure), and one individual that entered acclimation was not tested because of unexpected mortality, leaving the total number of individuals used in analysis at N=150.

## RESULTS

A total of 150 fish were successfully measured for SMR, of which 76 had the *vgll3*\*LL genotype (34 males and 42 females) and 74 had the *vgll3*\**EE* genotype (35 males and 39 females) (Appendix 2 - Table A3). Mean body length and mass was 0.95±0.03 g and 41.04±0.37 mm (±SE) for *vgll3*\**EE* fish and 1.2±0.03 g and 44.14±0.43 mm for *vgll3*\*LL fish (See note on body mass distributions in the statistical analysis section). *vgll3* genotype had no detectable effect on body condition (p=0.33, Appendix 2 - Table A2).

The *vgll3* genotype had no significant effect on SMR (Fig. 3). There was also no significant effect of sex (p=0.59) or the sex:*vgll3* interaction (p=0.69). Both the model excluding *vgll3* and the model excluding sex and *vgll3* as fixed factors had a lower AIC and Root-mean-square deviation (from cross-validation) than the full model, indicating that the models excluding *vgll3* and sex had the higher relative quality and higher predictive power. The metabolic scaling coefficient was estimated to 0.88±0.05 (95% CI) in the full model and 0.89±0.24 in the simple model, and the coefficient of variation was 6.23 and 6.19% for the simple and full model, respectively. The coefficient of variation was calculated as the SD of residuals after scaling them from log_10_ to log_e_ scale, as suggested by Garland (1984). See Appendix 2 - Table A1 for the full statistical summary.

**Figure 3.**
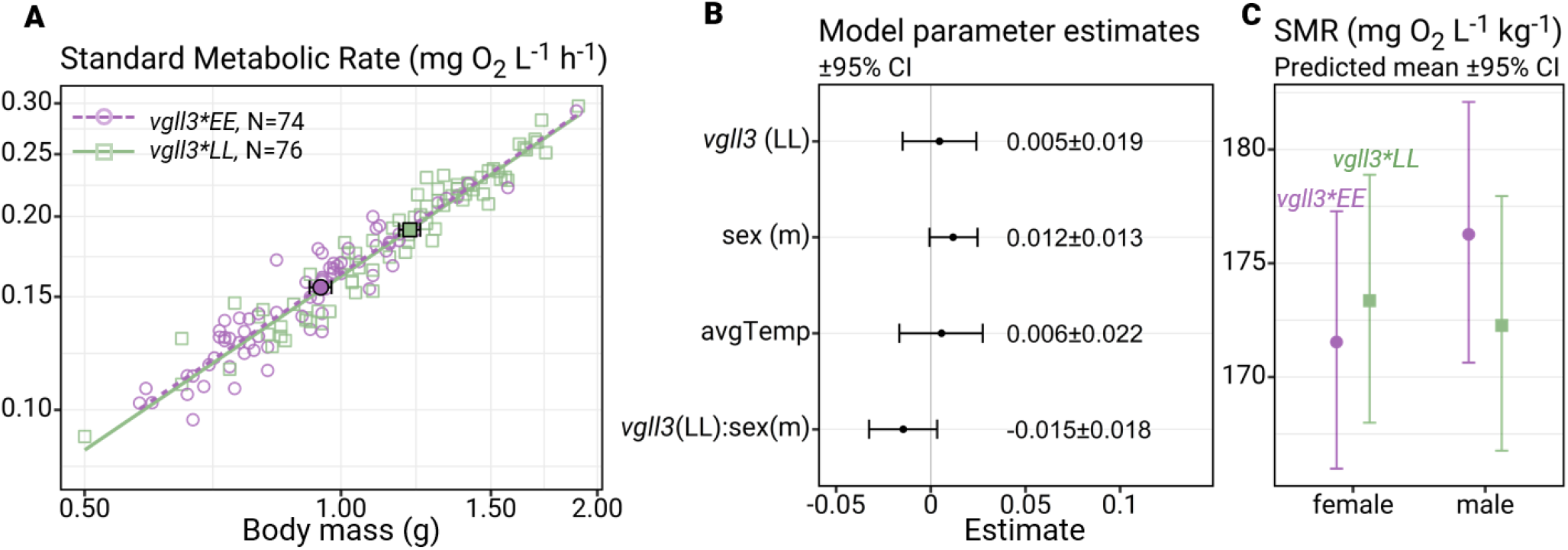
Main results. A) Log_10_-log_10_ plot of standard metabolic rate (mg O_2_ L^-1^ h^-1^) and body mass (g) for every included individual (N=150), with green squares indicating individuals with *vgll3*\**LL* genotype and purple circles indicated *vgll3*\**EE* genotype. Axis labels are back-transformed to show non-log values. The square and circle with a black border indicate genotype means, and horizontal error bars indicate the standard error for mean body mass. Note: An earlier experiment had sampled individuals from the study-families non-randomly in relation to size, resulting in a significant body-size difference between *vgll3* genotypes. B) Estimates of model coefficients (with 95% confidence intervals) for a mixed linear model of log_10_ standard metabolic rate, where we used log_10_ body mass, *vgll3* genotype, sex, vgll3:sex interaction, and average temperature of the rearing tanks before acclimation period as fixed effects, and family and respirometry batch as random effects. The estimates of the intercept (−0.85±0.570, 95%CI) as well as the parameter for log_10_ body mass (0.88±0.05, 95%CI) are not shown. The full model summary is shown in Appendix 2 - Table A1. C) Predicted means (with 95% confidence intervals) of mass-specific standard metabolic rate for male and female individuals of either *vgll3*\**EE* or *vgll3*\**LL* genotypes, fixed for a body mass of 1.08 g and an average temperature of 10.28°C (preceding the respirometry acclimation and trial) at the population level (no family effects), based on the model estimates in Fig. 3B.

Using the residual standard deviation of 0.027 obtained from the full model, a post hoc power analysis was performed to estimate the minimum effect size this study should have been able to detect with a power of 80%. Repeatedly simulating datasets (Appendix 2 - Fig. A2) where SMR is affected by *vgll3* genotype and sex as specified in the full model – and testing different values for the effect of *vgll3* and the *vgll3*:sex interaction – we found a minimum effect size of 4.25% and 6% on mass-adjusted (non-log) SMR for the *vgll3* and the *vgll3*:sex effects, respectively (translating to 0.018 and 0.025 for the model estimates). The power analysis indicates that although we did not detect a *vgll3* effect in this experiment, we cannot rule out effects of *vgll3* and *vgll3*:sex on mass-adjusted SMR smaller than 4.25% and 6%.

## DISCUSSION

*Vgll3* genotype has been shown to have a strong effect on salmon life-history (Ayllon et al., 2015; Barson et al., 2015; Verta et al., 2020), and recent findings have suggested that this effect might be mediated by *vgll3*-induced differences in body condition (Debes et al., 2021). Taking this together with the connections between standard metabolic rate (SMR) and resource acquisition and assimilation (Armstrong et al., 2011; Auer, Bassar, et al., 2020; Auer, Solowey, et al., 2020; Bochdansky et al., 2005; Millidine et al., 2009; Rosenfeld et al., 2015), we hypothesized that *vgll3* genotype might be asserting some of its effects through changes in energetic physiology that would be reflected in SMR. However, we did not detect any difference in either SMR or body condition between juvenile Atlantic salmon individuals of different homozygous *vgll3* genotypes. The lack of difference in body condition might be related to body size or life-stage; the fish in the study by Debes et al. (2021) were considerably more developmentally advanced, averaging 19 g, and had been reared for approximately 3100 degree days at stable daily temperatures with many of the males maturing the same autumn. In comparison, the fish in this study averaged just 1.08 g and had been reared at approximately 1900 degree days and also under a variable temperature regime, which could have further reduced growth (Imholt et al., 2011). It might then be possible that the body-condition-effects of *vgll3* genotype only start to occur at a larger body mass, or in the months prior to maturation. Considering the SMR results, measurement error is one factor that could potentially mask *vgll3* effects, but the SMR measurements in this study were generally precise, with a metabolic scaling coefficient well within the range for what has been observed in other fishes (Clarke & Johnston, 1999) and a low coefficient of variation for the model’s residuals, indicating good precision of the results. Our post hoc power simulation based on our observed residual variation indicated that this experiment, with its comparably high number of individuals, should have been able to detect effects of *vgll3* genotype down to an effect size as small as 4.25% on mass-adjusted SMR, as well as a sex-by-*vgll3* genotype interaction effect down to 6% (Appendix 2 - Fig. A2). This would be a comparably weak effect considering that mass- and sex-adjusted metabolic rates among individuals have been reported to vary 2-3 fold within a population (Auer et al., 2018; Burton et al., 2011; Metcalfe et al., 2016). Given this, the lack of difference in SMR prior to maturation or prior to a change in body condition does not support SMR of juvenile salmon as a mediator of *vgll3*-effects on age at maturity at this particular age and in these conditions. Finally, these results were supported by a parallel study using a different crossing design and juvenile salmon of larger body size (Prokkola et al., 2021) where no association between SMR and *vgll3* genotype was found.

The results indicate that any potential organismal differences caused by *vgll3* genotype do not affect physiological function or body composition in a way that changes SMR at this life stage in these conditions. Such differences could for example have been organ size (Rosenfeld et al., 2015), enzyme activity (Norin & Malte, 2012), mitochondrial leak respiration (Salin et al., 2016, 2019) or capacities related to digestion or resource acquisition/assimilation (Auer et al., 2015; Millidine et al., 2009). Alternatively, but less parsimoniously, there may be tissue-dependent variation in *vgll3*-effects on metabolic rates, but that these cancel each other out at the organismal level, resulting in similar whole-animal metabolic rate. For example, *vgll3*\**EE* individuals could have more metabolically active digestive systems (allowing for faster nutrient assimilation) or increased muscle mass (high MR), but then a higher investment in adipose tissue (low MR) masks this difference’s effect on the mass-specific metabolic rate. Nevertheless, our results indicate that energy-allocation effects of *vgll3* are not driven by transient whole-animal maintenance energy requirements at this particular life stage and in these conditions.

Life-history traits are tightly linked with fitness, and recent changes in the maturation timing of Atlantic salmon populations towards earlier maturation have been associated with an increased frequency of the *vgll3*\**E* allele (Czorlich et al., 2018). The lack of association observed here between *vgll3* genotype and SMR indicates that at this life stage, SMR will not necessarily be co-selected together with *vgll3* genotype. For example, SMR has been shown to associate with microhabitat preference in Atlantic salmon, where high-SMR individuals are better able to make use of the higher food availability found in faster-flowing microhabitats (Auer, Bassar, et al., 2020). Our findings indicate that selection on *vgll3* does not seem to constrain variation in SMR, and possibly by extension, might not constrain habitat use, behaviour or other correlated traits.

Metabolic phenotype is a multi-faceted set of traits, covering not just standard metabolic rate, but also the maximum metabolic rate, aerobic scope, and daily energy expenditure, to mention some. Additionally, the consequences of variation in metabolic traits are context-dependent and may change under different environmental conditions and life stages (Auer, Bassar, et al., 2020; Auer, Solowey, et al., 2020; Bochdansky et al., 2005; Millidine et al., 2009; Norin & Metcalfe, 2019). We initially planned to also measure metabolic rate under exhaustive swimming to obtain data on maximum metabolic rate (MMR), but had to abandon these trials as we were unable to motivate juvenile salmon of this age and size to do any exhaustive swimming, either by swim tunnel or by hand chasing. This behaviour might be specific to this life stage, as we were eventually successfully able to exercise fish up to MMR by hand chasing in a second experiment using a cohort from the same year that was studied 1–2 summer months later when the fish had grown from 1 g to 4 g (Prokkola et al., 2021). Besides supporting our finding on the lack of an association between SMR and *vgll3* genotype, Prokkola et al. (2021) found a significantly higher MMR and aerobic scope in vgll3*EE compared to *vgll3*\*LL individuals, indicating that *vgll3* genotype does affect the metabolic phenotype, but does so by affecting traits related to MMR without affecting SMR. In the wild, juvenile Atlantic salmon prefer staying in slow-velocity microhabitats where they act as sit-and-wait predators, darting out to catch suitable prey items as they pass by (Fraser et al., 1993; Metcalfe et al., 1997). It is thus possible that individuals at the life stage studied here are not well physiologically adapted to aerobically exhaustive swimming, but rather shorter bursts, due to their small relative muscle mass and glycogen stores available for exercise. The available literature reporting successful swimming respirometry or MMR measurements of salmon of ∼1g size is very sparse, and the experiments reported by (Dabrowski, 1986) and (Cutts et al., 2002) are to our knowledge the only ones that have been successful in motivating juvenile salmon close to or below this size to swim. We therefore recommend focusing efforts on investigating MMR at the latter end of the early life stage, or carefully devising alternative ways of inducing MMR.

## Conclusions

We found that the *vgll3* age-at-maturity genotype did not significantly affect standard metabolic rate in Atlantic salmon juveniles. Our results indicate that *vgll3’s* effect on age at maturity and resource allocation is unlikely to be mediated through maintenance energy requirements or related traits and that juvenile individuals of different *vgll3* genotypes face similar maintenance energy requirements.

## Author contributions

Conceptualization: CRP, JMP, ERÅ, SM, TA. Methodology: JMP, ERÅ, SM, CRP, TA Formal analysis: ERÅ, SM. Investigation: ERÅ, JMP, SM. Resources: CRP, TA. Data curation: ERÅ, SM, CRP. Writing – original draft: ERÅ, JMP, CRP. Writing – review & editing: ERÅ, JMP, CRP, SM, TA. Software: SM. Validation: ERÅ, JMP, SM, CRP. Visualization: ERÅ, JMP. Supervision: JMP, CRP. Project Administration: CRP, JMP, ERÅ. Funding Acquisition: CRP, TA, ERÅ.

## Acknowledgements

We thank Petra Liljeström, Paul Bangura, Katja Maamela, Markus Lauha, Suvi Ikonen, Nikolai Piavchenko, Anna Toikkainen and Seija Tillanen for their work with animal husbandry and system maintenance; Heidrikur Bergsson for designing the 3D-printed chamber caps; Markus Haapala for 3D-printing the chamber caps; Nordic University Hub NordicPOP (Nordforsk, project no. 85352) for providing access to the 3D printer; Annukka Ruokolainen and Shadi Jansouz for genotyping and sexing fin-clips; Tommy Norin for valuable input and advice on building the respirometry setup; Lammi Biological Station for their flexibility in enabling us to conduct this experiment despite the ongoing COVID pandemic; Anna Toikkanen, Antti Miettinen, Jacqueline Moustakas-Verho, Jukka-Pekka Verta, Marion Sinclair-Waters, Nikolai Piavchenko, Shadi Jansouz, Teemu Mäkinen & Laukaa hatchery staff for help with egg and milt collection and/or fertilizations; and Natural Resources Institute Finland (LUKE) for access to the broodstock.

## Data accessibility

The full datasets and the R scripts used to analyse them are available in Zenodo. DOI:10.5281/zenodo.5255061

## Appendix 1 Supplementary Materials and methods

### Respirometer design

Respirometers (Fig. A1) were designed for intermittent-flow respirometry (Forstner, 1983; Steffensen et al., 1984) with optical oxygen probes. The main chamber of the respirometers consisted of a glass tube (Length 70 mm, inner diam. 24 mm) (Schott, Finnish Special Glass, Espoo, Finland) with 3D-printed ‘HeiBer’ plastic caps (Heidrikur, 2020) sealed using a nitrile rubber o-ring at each end. Each chamber cap had two outlets, one for circulatory flow and one for flush flow. Inside the chamber, a flat circular 3D-printed plastic baffle partly covered each chamber cap to protect the outlets and to ensure a mixed water flow. The baffles were secured to the chamber caps using a stainless steel screw. The chamber caps and baffles were 3D printed using polyethylene terephthalate glycol (PETG, Devil Design, Mikolow, Poland)

Respirometer volume was measured using the weight difference between dry respirometers and respirometers filled with water (while plugging the flush-flow and oxygen probe outlets and inlets). Respirometers had a total volume of 47.3 ml, and the fish had an average weight of 1.08±0.3g (±SE) giving a 2.2% ratio of animal body mass to volume of respirometer.

For each chamber, one 3-6V DC pump acted as a circulation pump (unbranded, China) and was connected with Tygon tubing (Tygon S3 E-3603, Saint-Gobain, Paris, France) to one outlet on each chamber cap, providing a circulatory flow through the chamber. A 5-12V DC pump (DC30C, Anself, China) acting as a flush pump was connected to one chamber cap, and a Tygon tube reaching above the water surface was connected to the opposite chamber cap, serving as an overflow outlet during flushing. Flow speed was set to 0.7cm s^-1^ during the closed phase and 1.8 cm s^-1^ during the flush phase. Fish were able to lie still on the bottom of the chamber during both phases.

The oxygen probe was connected in-line with the circulation circuit using a plastic t-junction piece, sealed by threading the probe through a thin silicone tube before inserting it into the t-piece.

The 16 respirometers were divided into groups of four, wherein each group one 4-channel oxygen logger (FSO2-C4, Pyroscience, Aachen, Germany) recorded the oxygen concentrations from the oxygen probes (OXROB10, Pyroscience, Aachen, Germany), and one PumpResp pump control unit (4-channel model, FishResp, University of Helsinki, Finland, https://fishresp.org/pumpresp) automated the work of the flush and circulation pumps. Each oxygen logger used one PT100 temperature probe (TSUB21-CL5, Pyroscience, Aachen, Germany) to record the temperature of the respirometry tank near its respective respirometers. The oxygen loggers were set to automatically temperature-compensate the recorded oxygen concentrations. A single pump provided flush flow to all four respirometers. Oxygen probes were calibrated in O_2_-free water (deoxygenated using sodium sulphate) and air-saturated water.

**Figure A1.**
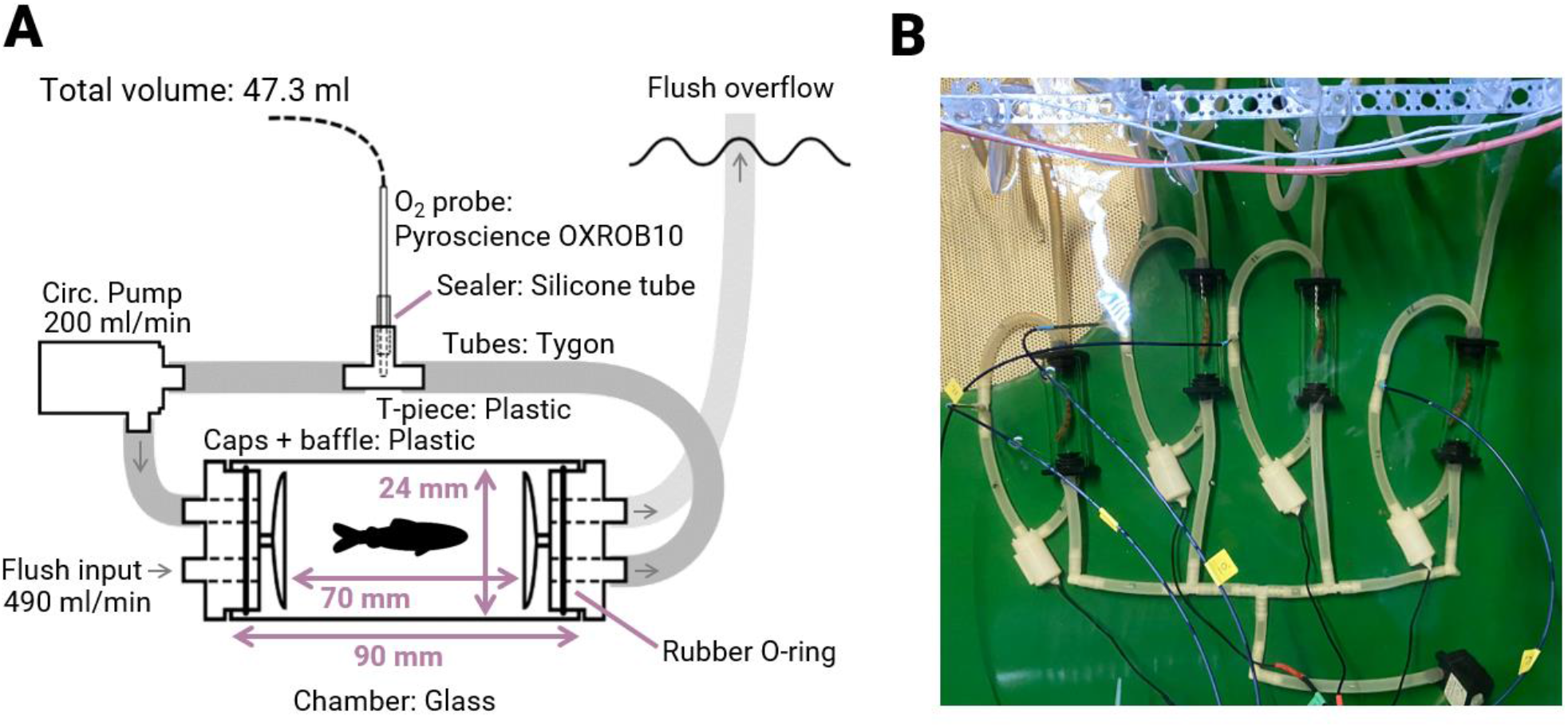
Design of respirometers. A) Schematic (not to scale) overview of a single respirometer. B) One group of four respirometers, in the respirometry tank, with fish, shown from above.

### Respirometry holding system

The full respirometry system consisted of three square 164 L tanks, two acclimation tanks and one respirometry tank, all located in a different room from the holding tanks. As opposed to the holding tanks, which had variable temperatures depending on the temperature of the incoming lake water, the respirometry system was set up to use the same water but to hold a constant temperature of 10.5°C. This was achieved by setting up two 8°C cold-water reservoir tanks, which both received an incoming 1 L min^-1^ flow of filtered (UV and a 60 µm physical filter) lake water, but were cooled by two separate aquarium coolers. Each of the respirometry system’s three tanks had a temperature-controlled pump which created a slow flow of 8°C water from the reservoirs into the tank whenever the tank temperature got higher than 10.5°C, achieving a highly stable temperature at 10.53±0.02°C (mean±SD) for the entirety of the experiment. An overflow pipe drained water from the acclimation tanks back to the cold-water reservoir. The two acclimation tanks shared one cold-water reservoir while the respirometry tank used the other. Both the respirometry tank and its cold -water reservoir were continuously UV filtered to reduce the growth of algae and background respiration. Water in all three tanks was continuously aerated with air-stones, and each acclimation tank had two Eheim 300 pumps (Eheim, Deizisau, Germany) to keep water mixed. Lighting was provided by one 80-cm Juwel Novolux LED-lights (JUWEL Aquarium, Rotenburg, Germany) for each acclimation tank, with photoperiod adjusted according to the local latitude.

### Extraction of respirometry data

All oxygen traces were inspected visually for linearity (and blindly from *vgll3* genotype), using regression diagnostic plots (i.e., residuals vs fitted, normal Q-Q, and scale-location), and 1423 slopes were excluded, leaving an average of 47.1±5.1 (±SD) MO_2_ measurements per individual used in the statistical analysis. Using the r^2^ of the fitted traces to inspect for linearity was dropped as only 0.6% of all slopes (54 out of 8739) failed to pass the coefficient of determination threshold (r^2^< 0.95), despite spontaneous fish activity and occasional malfunction of circulation pumps. MO_2_ was calculated from each of these slopes as described by Morozov et al. (2020) using the R package FishResp. Mean r^2^ for all oxygen traces used in the statistical analysis was 0.997±0.004 (±SD, N=7311). For each individual, the standard metabolic rate (SMR) was defined as the mean of the lowest normal distribution after having fitted all MO_2_ measurements to up to four normal distributions as described by Chabot et al. (2016). This method automatically separates the MO_2_ measurements into distributions representing the acclimation time, the measurements where the fish are quiescent, and measurements following potential spontaneous activity. In FishResp (Morozov et al. 2020), background respiration was modelled to increase linearly (based on pilot tests) from the pre-test measurement to the post-test measurement, and oxygen consumption was controlled for background respiration by subtracting the background respiration (and its predicted linear increase) from the oxygen concentration measurements before calculating the rate of oxygen consumed by the fish.

## Appendix 2 Supplementary analysis and data

**Figure A2.**
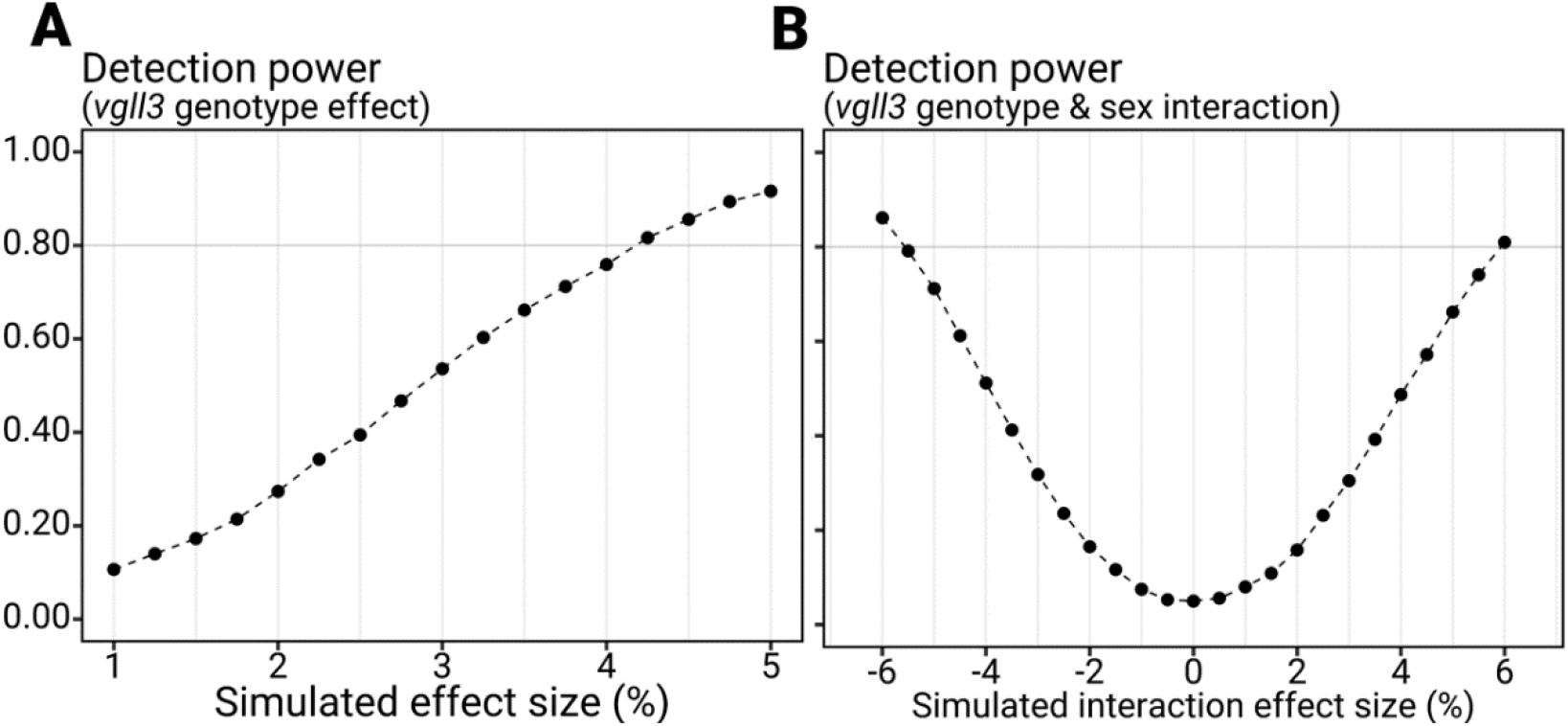
Power analysis. Shows the predicted power for this study (N=150 individuals, alpha=0.05, residual sd=0.027) to detect an effect on SMR from *vgll3* genotype and an interaction between *vgll3* genotype and sex, given different effect sizes. The analysis gives an indication of the smallest effect sizes this study can detect, given that *vgll3* genotype and sex affects SMR as in the model written below. Effect sizes are shown as the degree that the factors affect the mass-adjusted metabolic rate (mg O_2_ L^-1^h^-1^kg^-1^). For example, a 4% effect size would increase or decrease mass-specific metabolic rate with 4%. Each point represents the proportion of 10000 simulated datasets where a significant effect was found. Simulated datasets are based on a model of the form: *log(SMR) = a + b ×log(body mass) + c×vgll3EE + d×vgll3EE×male + e*, where a is the intercept; b is the metabolic scaling coefficient, set to 0.8; c and d are the effect sizes of genotype and the interaction between genotype and sex, respectively; and e is the error component, which here is normally distributed with a mean of 0 and a standard deviation of 0.027 (taken from results of this study). In the simulation for effect of genotype, the genotype-sex interaction was fixed at 0%. In the simulation for interaction of genotype and sex, the effect of genotype was fixed at 4%. The simulated datasets used body mass sampled randomly from a normal distribution with a mean of 1.08 g and a standard deviation of 0.3 (taken from results of this study), balancing *vgll3* genotype and sex evenly across the 150 simulated individuals. The analysis is available in the uploaded R script (see data accessibility).

**Table A1.** Statistical summaries for mixed-effect models on log_10_ standard metabolic rate. F and p values are calculated using Satterthwaite’s method using type III (full model) or type II (simple model) tests. Mixed models are fitted using restricted maximum likelihood. A) Summary for the full model, including the fixed effect of sex (m/f) and *vgll3* genotype (EE/LL), as well as their interaction. B) Summary for the simple model, excluding fixed effect of sex or *vgll3* genotype. C) Model comparison between full model, simple model, and a sex-only model (excluding *vgll3* but keeping sex as fixed factor). The Root-mean-square error (RMSE) is the result of a cross-validation of the models, which is obtained by splitting the dataset into two sets of odd- and even-numbered respirometry batches, and using model parameters estimated from one set to predict the results of the other set. The RMSE represents the deviation between the predictions and the real data, and lower values indicate a higher predictive ability. Batch was excluded as a random effect from the models in the cross-validations due to singularity. The Akaike information criterion (AIC) is an estimation of the relative quality of the models, taking residual variation and model complexity into account; A lower value indicates higher relative quality.

| Fixed effects | Estimate | 2.5% CI | 97.5% CI | Sum Sq | Mean Sq | DenDF | F<br>(Type III) | p |
| --- | --- | --- | --- | --- | --- | --- | --- | --- |
| Intercept | -0.853 | -1.077 | -0.062 |  |  |  |  |  |
| Log<sub>10</sub> Body mass | 0.878 | 0.830 | 0.928 | 0.844 | 0.844 | 90.347 | 1160.548 | >0.001 |
| *vgll3* (LL) | 0.004 | -0.014 | 0.024 | 0.000 | 0.000 | 12.943 | 0.087 | 0.771 |
| Sex (male) | 0.012 | -0.001 | 0.025 | 0.0007 | 0.0007 | 129.139 | 0.984 | 0.322 |
| AvgTemp | 0.006 | -0.017 | 0.027 | 0.0002 | 0.0002 | 6.831 | 0.228 | 0.647 |
| *vgll3*(LL):Sex(male) | -0.014 | -0.033 | 0.003 | 0.002 | 0.002 | 136.132 | 0.248 | 0.117 |
| Random effects | Std.dev |  |  |  |  |  |  |  |
| Family | 0.014 |  |  |  |  |  |  |  |
| Batch | 0.003 |  |  |  |  |  |  |  |
| Residual | 0.0269 |  |  |  |  |  |  |  |

**B) Simple model:** Log<sub>10</sub>smr ~ Log<sub>10</sub>mass + AvgTemp + (1|family) + (1|batch)
| Fixed effects | Estimate | 2.5% CI | 97.5% CI | Sum Sq | Mean Sq | DenDF | F<br>(Type II) | p |
| --- | --- | --- | --- | --- | --- | --- | --- | --- |
| Intercept | -0.833 | -1.072 | -0.590 |  |  |  |  |  |
| Log<sub>10</sub> Bodymass | 0.885 | 0.837 | 0.932 | 0.968 | 0.968 | 90.954 | 1318.099 | >0.001 |
| AvgTemp | 0.004 | -0.019 | 0.027 | 0.0001 | 0.0001 | 6.744 | 0.131 | 0.729 |
| Random effects | Std.dev |  |  |  |  |  |  |  |
| Family | 0.014 |  |  |  |  |  |  |  |
| Batch | 0.0002 |  |  |  |  |  |  |  |
| Residual | 0.0271 |  |  |  |  |  |  |  |

**C) Model Comparisons**
|  | RMSE | AIC |
| --- | --- | --- |
| **Full model:** |  |  |
| Log<sub>10</sub>smr ~ Log<sub>10</sub>mass + *vgll3* + sex + *vgll3*: sex + AvgTemp + (1\|family) + (1\|batch) | 0.0646 | -580.185 |
| **Sex only:** |  |  |
| Log<sub>10</sub>smr ~ Log<sub>10</sub>mass + sex + AvgTemp + (1\|family) + (1\|batch) | 0.0653 | -596.734 |
| **Simple model:** |  |  |
| Log<sub>10</sub>smr ~ Log<sub>10</sub>mass + AvgTemp + (1\|family) + (1\|batch) | 0.0656 | -606.652 |

**Table A2.** Statistical summaries for mixed-effect models checking for differences in body condition between *vgll3* genotypes. Body condition is calculated as Fulton’s K, where the body condition is the body mass (g) divided by the cube of the body length (mm) and multiplied by 100. F and p values are calculated using Satterthwaite’s method using type III (full model) or type II (sex-only model) tests. Mixed models are fitted using restricted maximum likelihood. A) Summary for the full model, including fixed effect of sex (m/f) and *vgll3* genotype (EE/LL), as well as their interaction. B) Summary for the sex-only model, excluding effect of *vgll3* genotype. C) Model comparison between full model, and sex-only model. The Root-mean-square error (RMSE) is the result of a cross-validation of the models, which is obtained by splitting the dataset into two sets of odd- and even-numbered respirometry batches, and using model parameters estimated from one set to predict the results of the other set. The RMSE represents the deviation between the predictions and the real data, and lower values indicate a higher predictive ability. Batch was excluded as a random effect from the models in the cross-validations due to singularity. The Akaike information criterion (AIC) is an estimation of the relative quality of the models, taking residual variation and model complexity into account; A lower value indicates higher relative quality.

| Fixed effects | Estimate | 2.5% CI | 97.5% CI | Sum Sq | Mean Sq | DenDF | F<br>(Type III) | p |
| --- | --- | --- | --- | --- | --- | --- | --- | --- |
| Intercept | 1.353 | 1.315 | 1.390 |  |  |  |  |  |
| *vgll3*(LL) | 0.016 | -0.035 | 0.065 | 0.004 | 0.004 | 12.177 | 1.049 | 0.326 |
| Sex (male) | -0.018 | -0.045 | 0.010 | 0.003 | 0.003 | 136.196 | 0.785 | 0.377 |
| *vgll3*(LL):Sex(male) | 0.018 | -0.023 | 0.058 | 0.003 | 0.003 | 130.933 | 0.725 | 0.396 |
| Random effects | Std.dev |  |  |  |  |  |  |  |
| Family | 0.040 |  |  |  |  |  |  |  |
| Batch | 0.020 |  |  |  |  |  |  |  |
| Residual | 0.061 |  |  |  |  |  |  |  |

**B) sex-only model:** condition ~ sex + (1|family) + (1|batch)
| Fixed effects | Estimate | 2.5% CI | 97.5% CI | Sum Sq | Mean Sq | DenDF | F<br>(Type II) | p |
| --- | --- | --- | --- | --- | --- | --- | --- | --- |
| Intercept | 1.360 | 1.331 | 1.389 |  |  |  |  |  |
| Sex (male) | -0.009 | -0.031 | 0.011 | 0.003 | 0.003 | 137.06 | 0.844 | 0.360 |
| Random effects | Std.dev |  |  |  |  |  |  |  |
| Family | 0.040 |  |  |  |  |  |  |  |
| Batch | 0.0004 |  |  |  |  |  |  |  |
| Residual | 0.004 |  |  |  |  |  |  |  |

**C) Model Comparisons**
|  | RMSE | AIC |
| --- | --- | --- |
| **Full model:**<br>condition ~ *vgll3* + sex + *vgll3*: sex + (1\|family) + (1\|batch) | 0.073 | -345.384 |
| **Sex only:**<br>condition ~ sex + (1\|family) + (1\|batch) | 0.072 | -359.229 |

**Table A3.** Count of included individuals (N=150) for each *vgll3* genotype, family and sex.

| *vgll3\*EE* | Fam.2 | Fam.3 | Fam.7 | Fam.8 | Fam.9 | Fam.11 | Fam.13 | Total |
| --- | --- | --- | --- | --- | --- | --- | --- | --- |
| N Males | 6 | 7 | 3 | 5 | 5 | 5 | 4 | 35 |
| N Females | 5 | 4 | 6 | 4 | 6 | 6 | 8 | 39 |
| Total | 11 | 11 | 9 | 9 | 11 | 11 | 12 | 74 |

| *vgll3\*LL* | Fam.1 | Fam.4 | Fam.5 | Fam.6 | Fam.10 | Fam.12 | Fam.14 | Total |
| --- | --- | --- | --- | --- | --- | --- | --- | --- |
| N Males | 4 | 8 | 6 | 3 | 5 | 4 | 4 | 34 |
| N Females | 8 | 3 | 6 | 9 | 5 | 4 | 7 | 42 |
| Total | 12 | 11 | 12 | 12 | 10 | 8 | 11 | 76 |

